# Focused ultrasound blood brain barrier opening mediated delivery of MRI-visible albumin nanoclusters to the rat brain for localized drug delivery with temporal control

**DOI:** 10.1101/696237

**Authors:** Megan C. Rich, Jennifer Sherwood, Aundrea F. Bartley, Quentin A. Whitsitt, W.R. Willoughby, Lynn E. Dobrunz, Yuping Bao, Farah D. Lubin, Mark Bolding

**Author notes:** Megan C. Rich –, Aundrea F. Bartley –, W. R. Willoughby –, Lynn E. Dobrunz –, Yuping Bao –, Farah D. Lubin –. corresponding author, 1670 University Blvd, Volker Hall G082, Birmingham, AL, 35294.

## Abstract

There is an ongoing need for noninvasive tools to manipulate brain activity with molecular, spatial and temporal specificity. Here we have investigated the use of MRI-visible, albumin-based nanoclusters for noninvasive, localized and temporally specific drug delivery to the rat brain. We demonstrated that IV injected nanoclusters could be deposited into target brain regions via focused ultrasound facilitated blood brain barrier opening. We showed that nanocluster location could be confirmed *in vivo* with MRI. Additionally, following confirmation of nanocluster delivery, release of the nanocluster payload into brain tissue can be triggered by a second focused ultrasound treatment performed without circulating microbubbles. Release of glutamate from nanoclusters *in vivo* caused enhanced c-Fos expression, indicating that the loading capacity of the nanoclusters is sufficient to induce neuronal activation. This novel technique for noninvasive stereotactic drug delivery to the brain with temporal specificity could provide a new way to study brain circuits *in vivo* preclinically with high relevance for clinical translation.

**Graphical Abstract:** 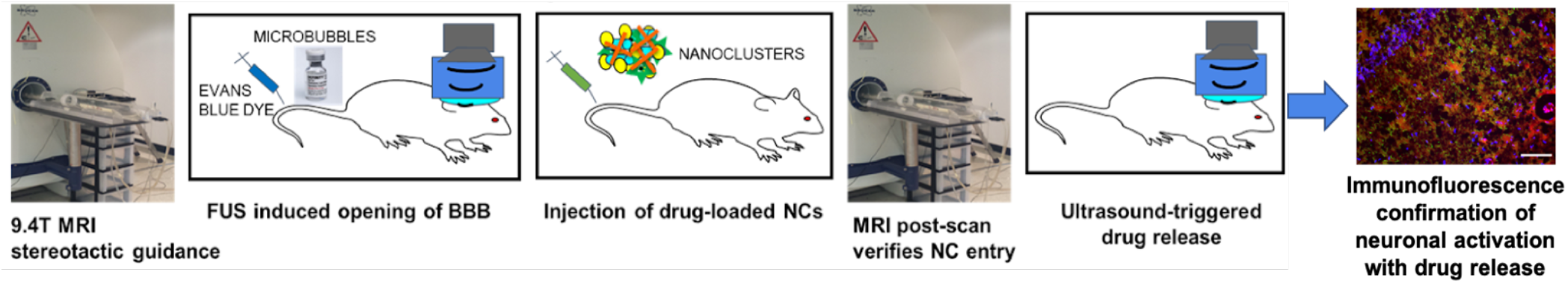

## Introduction

One major impediment to transforming basic neurobiological findings into treatments or therapies is the blood brain barrier (BBB) [1], which blocks passage of most drugs into the brain [2,3]. This prevents many neuroactive drugs from reaching their full potential as therapeutic agents or as research tools [4–6]. Of the drugs that cross the BBB, many have undesirable side effects, either in other brain regions or in the periphery, that can outweigh clinical benefit or confound preclinical experiments [4–6]. Existing methods of localized drug delivery to specific brain regions require invasive surgery to penetrate the skull and brain tissue for injections or cannula insertion [7]. The invasiveness of this surgery prevents its routine use in clinical settings and the tissue damage and subsequent inflammatory responses are ubiquitous confounds for preclinical studies [8]. The ability to noninvasively deliver drugs across the BBB and target them to specific brain regions will have tremendous impact on treatments for neurological disorders while simultaneously providing a powerful investigational tool for preclinical research that could be used in parallel, providing a tighter bench-to-bedside approach. One method of targeted drug transport across the BBB with minimal tissue damage is transcranial focused ultrasound (FUS) together with microbubbles to focally and transiently open the BBB [9–13]. FUS is an ideal tool for localized drug delivery due to its current use in humans [14,15] and its non-invasive nature [12,15,16]. However, brain localization of systemically delivered drugs with FUS BBB opening requires the use of drugs that do not readily cross the BBB. This limits which drugs can be delivered with this technique, including the majority of current clinical neurological agents. In addition, systemically circulating drugs can cause off target side effects. One strategy to overcome these limitations is albumin-based drug encapsulation which allows control over the location of drug delivery, prolongs circulation time and reduces off-target effects [17–19]. In addition, albumin based drug carriers can be targeted with ultrasound to induce release of their cargo [20,21], thus providing temporal control.

We have shown previously that albumin-based nanoclusters (NCs) can be embedded with ultrasmall iron oxide nanoparticles to provide MRI visibility and can encapsulate dye molecules as well as antibodies [19,22]. Furthermore, the NC load remained inside the NC unless digested with trypsin, and NCs remained stable *in vivo* when delivered by IV [19]. Here, we investigated whether these albumin-based NCs could be used in combination with MRI guided FUS BBB opening (BBBO) for localized drug delivery to the rat brain. We report that IV injected NCs can locally diffuse into the brain with FUS facilitated BBBO and provide MRI contrast at the delivery site. Once delivered to target brain regions, we were able to induce drug and dye release by a second FUS treatment, providing temporal control. We show that the NC load is sufficient to cause localized changes in neural activity following release of glutamate from FUS delivered NCs in vivo.

## Materials and methods

### Nanocluster formation and FUS induced drug release

NCs were created as described previously [19,22]. Briefly, first tannic acid-coated ultrasmall (<4 nm) iron oxide nanoparticles were synthesized [22] followed by crosslinking of nanoparticles with bovine serum albumin (BSA) proteins by the addition of ethanol and glutaraldehyde. The overall size of the NCs can be tuned by altering the amounts and rates of ethanol addition. Glutaraldehyde provides surface crosslinking of NCs, which subsequently inhibits further growth and aggregation of BSA NCs. For these studies, NCs between 100 nm and 200 nm were used at a concentration of 2 mg/mL.

### MRI procedure

Male Sprague Dawley rats weighing 250-350 g were obtained from Charles River Laboratories and housed in accordance with UAB Institutional Animal Care and Use Committee (IACUC) guidelines. Animals had free access to water and rat chow, and were maintained on a 12:12 hr light:dark cycle. Animals were anesthetized with isoflurane (1.5 – 3%) and positioned on a transportable 3D printed rat stereotaxic frame containing an MRI fiducial. Animals remained in the stereotaxic frame under isoflurane anesthesia throughout the MRI and FUS procedures. T_1_- and T_2_-weighted MR images were collected on a 9.4 T Bruker horizontal small-bore animal MRI scanner with a custom surface coil prior to FUS procedure (prescan), as well as 15 minutes, 30 minutes, and 1 hour after FUS BBBO. Image parameters were as follows, for T_1_- and T_2_-weighted axial images; width: 30 mm, height: 51.2 mm, depth: 3 mm, voxel size: 0.2× 0.2× 0.2 mm^3^, number of slices: 13; for T_1_-weighted coronal images; width: 30 mm, height: 30 mm, depth: 27 mm, voxel size: 0.2× 0.2× 1 mm^3^, number of slices: 27; for T_2_-weighted coronal images; width: 30 mm, height: 30 mm, depth: 27 mm, voxel size: 0.1× 0.1× 1 mm^3^, number of slices: 27. A pneumatic pillow sensor placed under the rat’s chest and connected through an ERT Control/Gating Module (SA Instruments) was used to monitor the rat’s respiratory cycle during imaging. During the prescan, hippocampal or anterior cingulate cortex coordinates were measured in 3 planes from the MRI fiducial which would later be used for FUS targeting (Figure 1b).

**Figure 1.**
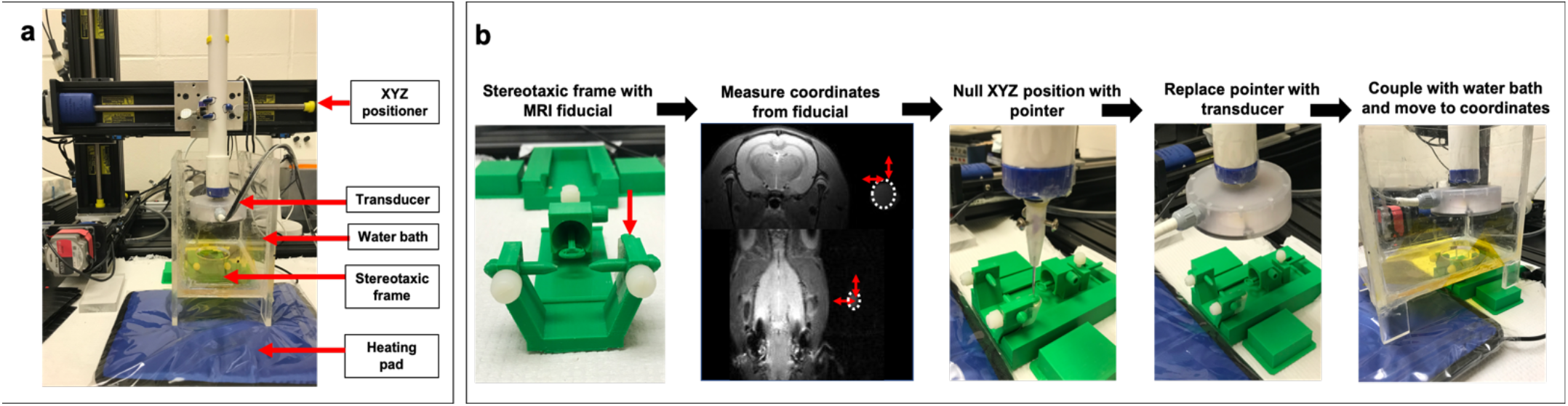
FUS setup and targeting procedure. (a) FUS benchtop setup. (b, from left to right)) Animals are first positioned in a 3-D printed stereotaxic frame equipped with an MRI fiducial (red arrow, left image). The frame is then placed inside the MRI bed and distance from the fiducial (dotted circle) to the target brain region is measured using both coronal images for the dorsal/ventral measurements and axial images for the rostral/caudal measurements, medial/lateral measurement can be collected from both axes. Animals are kept in the frame and transferred to the FUS station where a pointer is used to zero the XYZ positioner at the location of the fiducial. The pointer is then replaced with the FUS transducer (FUS Instruments) and a water bath is coupled to the animal’s head with gel. Lastly, using the XYZ positioner and the measurements collected from the MR images, the transducer focal point is positioned over the target brain site.

### FUS BBB opening and localized delivery of nanoclusters

Animals were transferred in the 3D printed stereotaxic frame from the MRI to the ultrasound equipment while maintained under anesthesia. The 3D printed stereotaxic frame slides into a 3D printed holder, thereby allowing the animal to remain in a static position under the ultrasound apparatus. Prior to FUS BBBO, a tail vein catheter was inserted and hair was shaved from the scalp and further removed with Nair. Using an XYZ positioning system (Velmex, Bloomfield, NY) driven by custom software written in LabVIEW (National Instruments, Austin TX), a pointer matching the transducer focal distance was positioned on the top and center of the MRI fiducial in the stereotaxic frame and this location was set to zero. The pointer was replaced with the ultrasound transducer and a water bath was coupled to the scalp with US gel and filled with degassed water. Using the coordinates that were determined during the MRI prescan, the transducer was moved to focus on the target brain region (Figure 1). Animals were then intravenously injected with 1 mL/kg of 3% Evans blue dye (EBD) (Sigma Aldrich), which was allowed to circulate for 5-minutes. Next, animals were then slowly infused (0.2 mL/2 mins) with 30 *μ*L/kg of Definity microbubbles (Lantheus Medical Imaging, Billerica, MA) in saline while FUS was applied to the target brain region. The FUS transducer was provided by FUS Instruments (Toronto, ON, Canada) and the FUS parameters were as follows: 0.30 MPa, 1.1 MHz, 10 ms burst, 1 Hz burst repetition rate, 2-minute duration. The Definity infusion and FUS exposure were repeated a second time following a 5-minute gap allowing for the first Definity injection to clear from circulation. Only one hemisphere was targeted with FUS BBBO, leaving the other hemisphere as an internal control. Immediately following FUS, animals were injected with either 2 mL/kg of unloaded NCs, NCs loaded with FITC dye, NCs loaded with glutamate (5 mM/mL, Sigma-Aldrich), 2 mL/kg of 5 mM/mL glutamate (Sigma-Aldrich) in saline, or 2 mL/kg of saline alone.

### Release-FUS procedure

Following the 30-minute MRI postscan, animals receiving release-FUS were removed from the MRI scanner and transported back to the ultrasound equipment while remaining in the stereotaxic frame. They were again placed into the stereotaxic frame holder, which provides the same positioning for the BBBO and release-FUS treatments. A water bath was placed over the scalp coupled with ultrasound gel and the FUS transducer was lowered back to its target coordinate. The animals were again treated with a 2-minute FUS exposure in the same target region using the same ultrasound parameters as stated above but without Definity microbubbles. FUS application in the absence of microbubbles does not cause the BBB to open further [23]. This was tested in several animals (data not shown), and it is evident in the similar degree of EBD staining between groups (Figures 4 and 5). In addition, release-FUS itself does not cause any c-Fos activation, as seen in our control animals (Figure 5).

### Nanocluster MRI contrast analysis

In order to analyze intrahippocampal NC contrast T_1_-weighted MRI images were first loaded into 3DSlicer software (https://www.slicer.org/) [25] for nonuniform intensity normalization using the publicly available N4ITK toolkit [26]. The images were then uploaded into FIJI [24], and using a rat brain atlas for reference, an ROI was drawn around both the FUS targeted and the non-targeted hippocampus. The minimum gray value from 3 ROIs per hemisphere were averaged and the ratio of non-target/target hemisphere values were analyzed.

### Brain tissue processing, immunofluorescence imaging and analysis

Following FUS BBBO and MR imaging, animals were perfused with 4% buffered formalin and brain tissue was immediately collected and frozen in Optimal Cutting Temperature compound (O.C.T., Tissue-Tek, Sakura Finetek USA). Brain tissue was stored at −80°C until cryostat sectioning. 10 *μ*m frozen sections were first thawed at room temperature for 10 minutes then fixed with 4% buffered formalin and washed 3 times with 1x PBS. Sections then underwent antigen retrieval by incubating in boiling sodium citrate solution, first for 2 minutes, then exchanged with fresh boiling solution and repeated twice for 5 minutes. Slides were then washed 3 times for 3 minutes in 1x PBS. Next, the sections were incubated in 5% goat serum block (Jackson Immunoresearch, West Grove, PA) for 1 hour at 4°C. Tissue sections were then incubated in primary antibody to either BSA (1:1000, ThermoFisher, Grand Island, NY), NeuN (1:400 MAB377, MilliporeSigma, St. Louis, MO) or c-Fos (1:1000, SC-52, Santa Cruz Biotechnology, Dallas, TX) overnight at 4°C followed by rinsing 3 times with 1x PBS. Sections were then incubated in secondary antibody (1:300, Alexa Fluor 488, or AffiniPure Mouse Anti-Rat IgG AMCA conjugate, Jackson Immunoresearch) for 2 hours at room temperature. Slides were then rinsed 3 times with 1x PBS and cover-slipped with either Fluoromount mounting medium (ThermoFisher, Grand Island, NY) or Vectashield Mounting Medium with DAPI Hardset (Vector Laboratories, burlingame, CA). Fluorescent microscopy imaging was performed using a Zeiss Axio-Imager microscope and processed using FIJI. Presence of FITC dye in tissue was quantified using an automated written macro for the performance of the particle analysis function in FIJI. For analysis of anti-c-Fos stained tissue, DAPI fluorescence images were used to get a total cell count using FIJI’s analyze particle function, which also generated outlines of cells which were overlaid onto the green channel depicting c-Fos fluorescence. c-Fos positive cells were counted using FIJI’s “point” tool. Cell counting was performed blinded to treatment condition, and only DAPI positive cell nuclei that overlapped with the green-c-Fos channel were counted. The percentage of total cells that were c-Fos positive were compared between groups.

## Results

### FUS BBB opening

We first tested the reliability of our FUS BBBO parameters and setup (specified above) by targeting both the anterior cingulate cortex (ACC) (n=5) and the hippocampus (n=9). In these experiments, in addition to EBD, the MRI contrast agent Gadovist, was IV injected (0.1 mL/ kg) immediately following FUS BBBO to verify targeted opening of the BBB. Figure 2a, c, f and g show enhanced MRI contrast where the Gadovist has entered the tissue through the opened BBB in vivo. BBBO was further confirmed by the spread of EBD which could be seen upon removal of the perfused brain both in the ACC (Figure 2b) and the hippocampus (Figure 2e). Note that the hippocampus is a deeper brain structure providing a fainter EBD appearance (Figure 2e) compared to the more superficial ACC target (Figure 2b). In addition, BBBO was evident by EBD fluorescence (excitation at 620nm, emission at 680nm) in 10 *μ*m brain sections from FUS treated animals (Figure 2, d and h).

**Figure 2:**
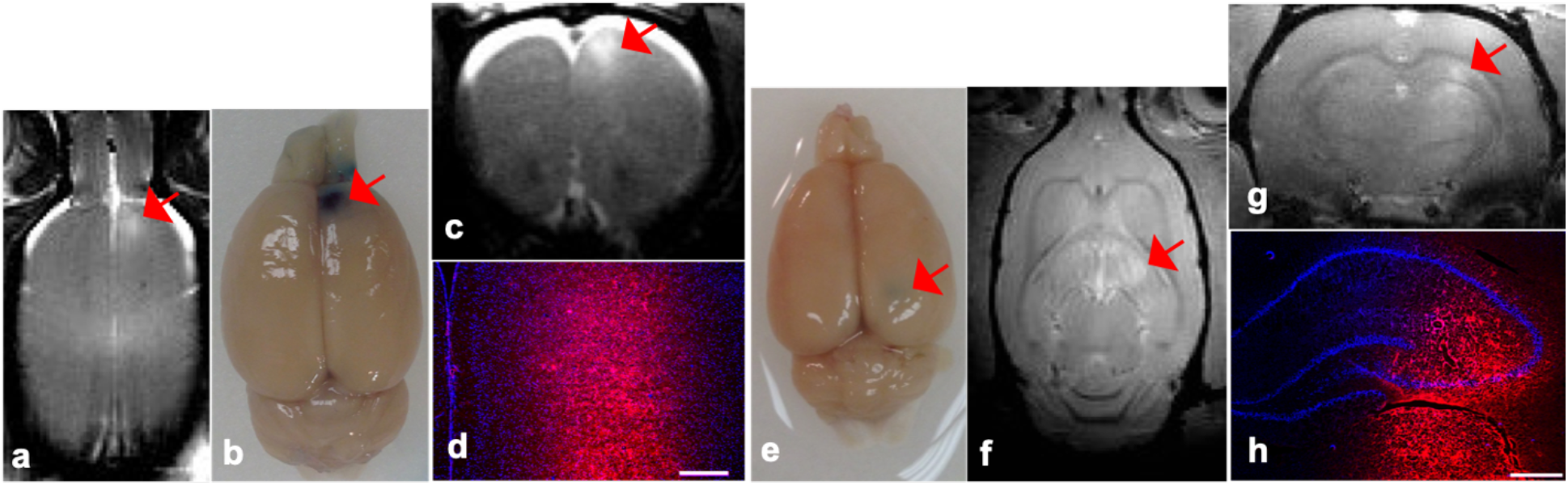
FUS BBB opening targeted to different locations in the rat brain. BBBO is evident by MRI contrast enhancement (a, c, f and g) and Evans blue dye (EBD) (b, d, e and h) targeted to the anterior cingulate cortex (a-d) and the hippocampus (e-h). FUS parameters: 0.3MPa, 1.1MHz, 10ms burst, 1Hz repetition rate, 120s duration, repeated 2x, simultaneous slow infusion of Definity. Scales bars: 500*μ*m (d and h).

### Localized nanocluster delivery with FUS induced BBB opening

To investigate whether NCs could be locally deposited into the hippocampus using FUS BBBO, animals were first pre-scanned by MRI at 9.4T, and then injected with EBD by tail vein catheter. FUS BBBO was then carried out, as described above, in the absence of injected Gadovist. Immediately afterwards, 2 mL/kg of HEPES buffer solution containing NCs (2 mg/mL) was injected into the rat by IV. Animals were then MR imaged again at 9.4T 15 minutes, 30 minutes and 1 hour after NC injection (Figure 3a). Control animals underwent the same BBBO procedure but received a saline injection without NCs. Animals treated with NCs showed enhanced MRI contrast at the FUS target site indicating NC entry into brain tissue with BBBO (Figure 3b). Hippocampal ROI analysis revealed that NC loading caused significant changes in MRI contrast at 15-minutes, 30-minutes and 1-hour post BBBO and NC injection (Figure 3c). Additionally, no significant contrast change was found in animals who received saline following BBBO (Figure 3c). Furthermore, NCs were identified in brain tissue by immunofluorescence staining of BSA (Figure 3d, bottom row) within areas of BBBO, which are shown by EBD fluorescence. No BSA immunoreactivity was observed in slices from animals with BBBO + saline injections (Figure 3d, top row). These results indicate that targeted and spatially-restricted entry of NCs into the brain can be achieved with FUS-induced BBBO.

**Figure 3.**
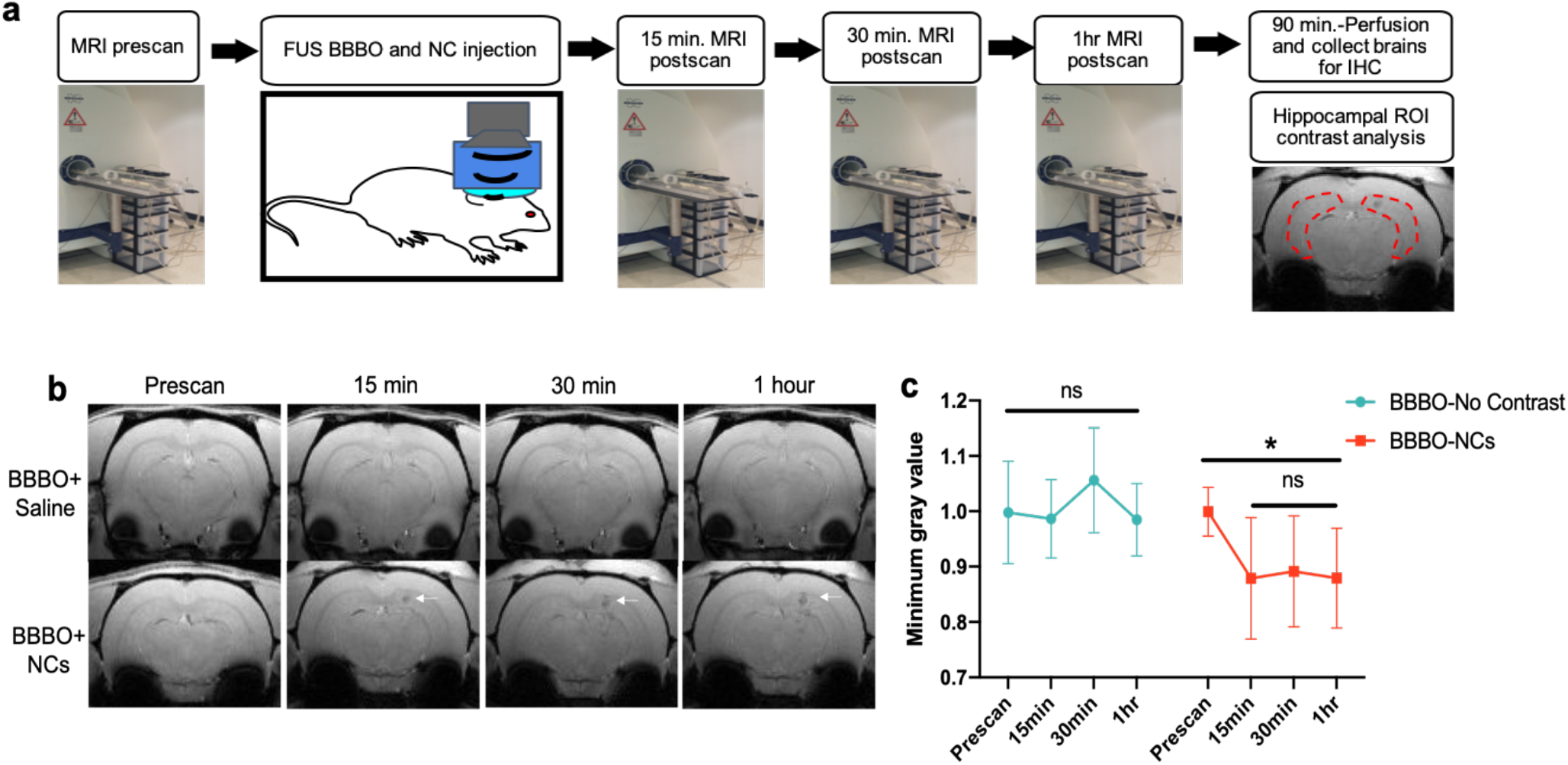
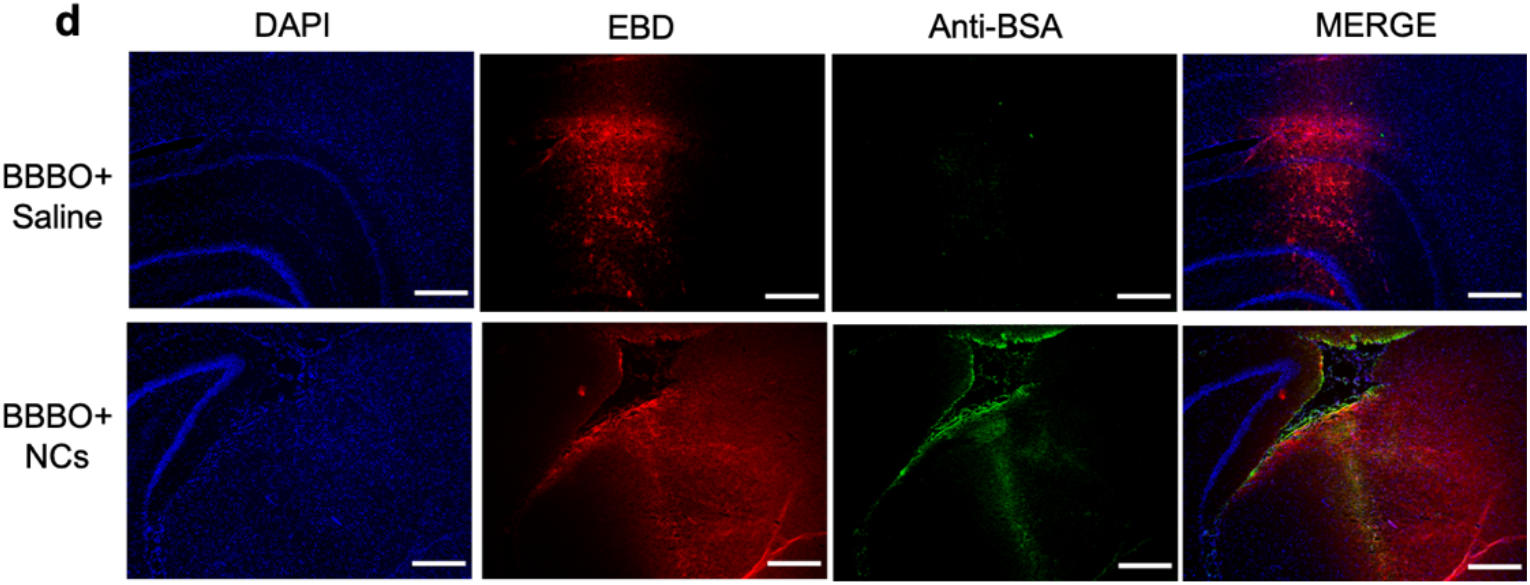
Nanoclusters load into brain tissue after localized BBB opening. (a) Procedure diagram. (b) T_1_-weighted MR images of an animal IV injected with saline (top) or NCs (bottom) immediately following BBBO in the hippocampus. NCs provide MRI contrast as they load into brain tissue at 15 min., 30 min. and 1hr after injection (white arrows). (c) Plotted MRI minimum gray values from hippocampal ROIs before and at 15 min, 30 min. and 1hr after FUS BBBO and NC (red) or saline (blue) injection (Time 0). Values are a ratio of target:non-target hippocampi and are presented as Mean±S.D., n=6-13 per group. All time points post NC delivery show significant changes in MRI contrast due to NC loading into brain tissue; statistical significance was set to *p* < .05 with \*\*\**p* < .001, ns=not-significant. (d) Representative immunofluorescence staining (n=3) with antibody to BSA showing NC distribution within BBBO region (EBD) in animals treated with unloaded-NCs (bottom) or saline (top) after BBBO. DAPI fluorescence shows overall cellular morphology. Scale bars; 100 *μ*m.

### FUS stimulated release of dye from nanoclusters

To test whether NCs could release their load *in vivo* with release-FUS, delivery of either FITC-loaded NCs or empty-NCs to one hemisphere of hippocampus was performed using FUS BBBO as described above. Animals were then exposed to a release-FUS treatment followed by perfusion and tissue collection (Figure 4a). Using fluorescent microscopy analysis of FITC dye in the region that was opened by FUS (EBD, Figure 4b), we observed a 500-fold increase in the amount of FITC dye in response to release-FUS (Figure 4b, c). Control animals that had empty-NCs or FITC-NCs without release-FUS exposure had little to no dye particles in the open BBB brain region (Figure 4b, c). These data suggest the successful delivery and release of NC-load into the target brain region.

**Figure 4:**
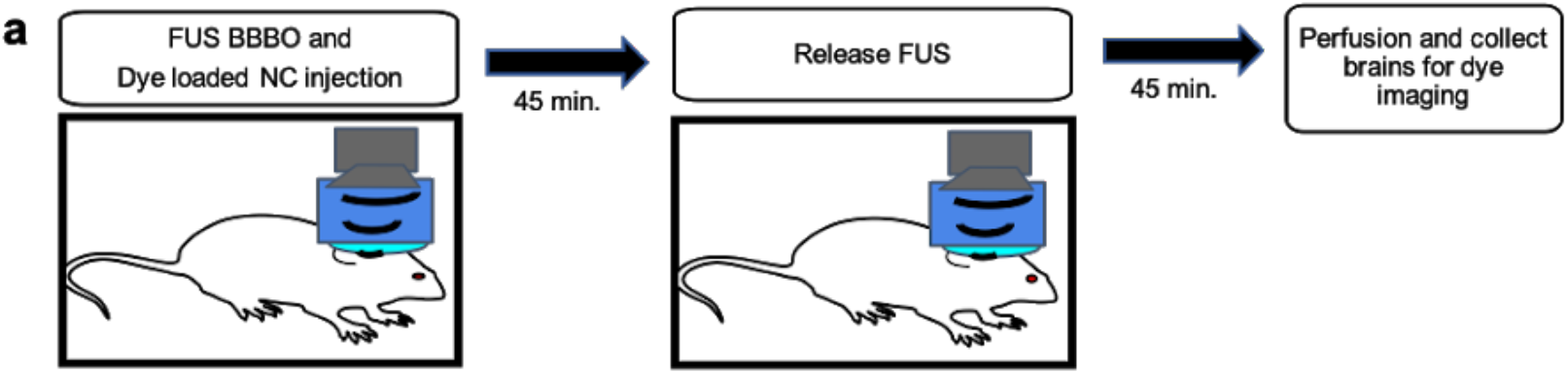
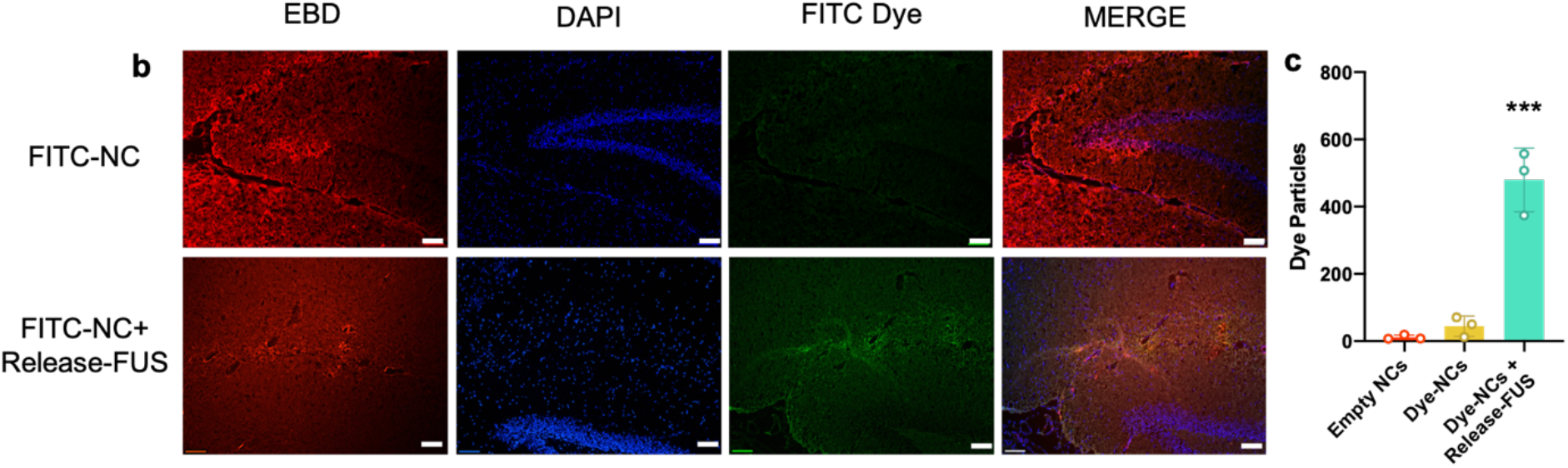
Dye release from NCs in vivo with release-FUS. (a) Experimental timeline; animals were injected IV with FITC-dye loaded NCs immediately after BBBO procedure. Animals were then exposed to a release-FUS treatment and brains were collected for fluorescence microscopy imaging. (b) Representative immunofluorescence images of hippocampi 90-minutes post BBBO and dye-loaded NC injection with (bottom row) and without (top row) a release-FUS treatment. EBD; Evans blue dye, indicating BBBO region, DAPI nuclear stain, indicating overall cellular morphology, FITC dye, indicating released dye from NCs. Scale bar: 100 *μ*m. (c) FITC particle quantification collected in brain sections from animals treated with empty NCs, Dye-loaded NCs and Dye-loaded NCs + release FUS. Values represent FITC particle number within a 1283 *μ*m X 958 *μ*m area of tissue. Statistical significance was set to *p* < .05 with \*\*\**p* < .0002.

### FUS stimulated release of glutamate from nanoclusters

In order to test whether drug loaded NCs could cause localized changes in brain activity in response to release-FUS, we delivered glutamate-loaded NCs (GluNCs) to the hippocampus *via* FUS BBBO followed by a release-FUS treatment (Figure 5a). In order to detect whether glutamate release from NCs caused neuronal activation, animals were sacrificed 90-minutes after release-FUS for anti-c-Fos immunofluorescence (Figure 5a). Animals that were treated with both GluNCs and release-FUS showed significantly enhanced c-Fos activation in the area of BBBO compared to animals that were treated with GluNCs without a release-FUS. These data suggest that glutamate was released from NCs at sufficient levels to activate neurons *in vivo* (Figure 5b, c). As a positive control, a separate group of animals were injected IV with free glutamate immediately after FUS BBBO and were sacrificed 90-minutes later. We observed that this group of animals had similar c-Fos levels to the group that received GluNC with release-FUS (Figure 5b, c). In addition, we confirmed that c-Fos levels were not significantly altered in tissue from animals who received FUS BBBO alone, FUS BBBO with empty NCs or FUS BBBO with release-FUS (Figure 5b, c), verifying lack of neural activation in these groups. This observation is consistent with previous studies, indicating that the applied FUS with the parameters in our studies does not modulate neural activity [27,28]. These data indicate that we were able to deliver and release enough glutamate into the brain to provide localized neuromodulation.

**Figure 5.**
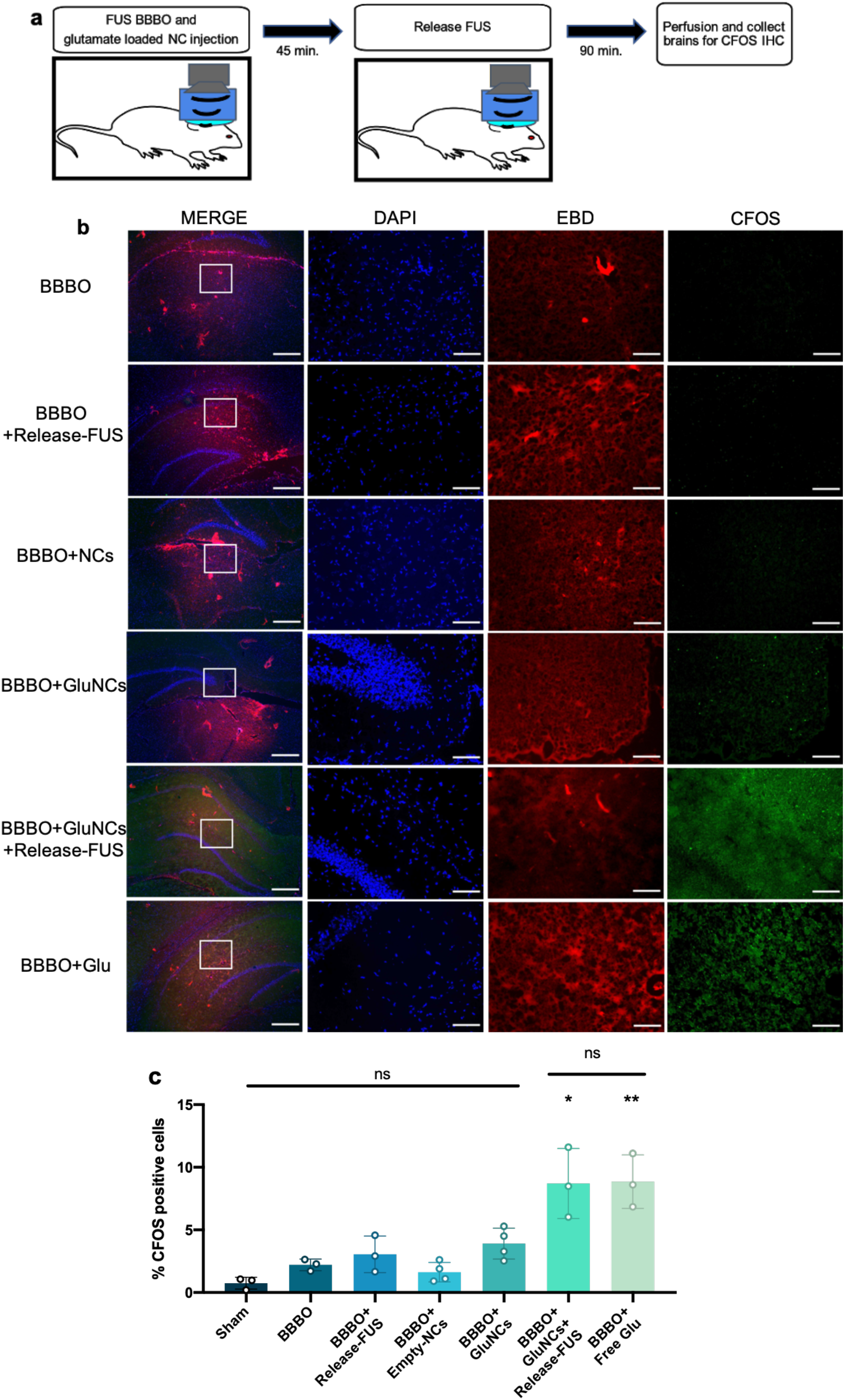
Release of glutamate from NCs in vivo causes neuronal activation. (a) Experimental timeline; animals were injected IV with glutamate loaded NCs immediately after the FUS BBBO procedure. Animals were then exposed to a release-FUS treatment and brains were collected 90 mins later for c-Fos immunofluorescence. (b) Immunofluorescence staining for c-Fos. Control groups included; Sham: no BBBO or injection, BBBO: BBBO only, BBBO+release-FUS, BBBO+empty NCs, BBBO+GluNCs: BBBO followed by glutamate loaded NC injection with no release-FUS treatment and BBBO+free Glu: animals were IV injected with 2 mL/kg of 5 mM/mL glutamate in saline immediately after BBBO procedure. Left column inlet represents the 5x higher magnification in adjacent images; MERGE column scale bar 500 *μ*m, adjacent images scale bar; 100 *μ*m. EBD; Evans blue dye, indicating BBBO region, DAPI nuclear stain, indicating overall cellular morphology, c-Fos indicating neuronal activation. (c) Percent c-Fos positive cells were significantly higher in animals who received GluNCs with a release-FUS treatment and in animals who received free glutamate compared to all other groups. Values indicate average percent c-Fos positive cells within 641 *μ*m × 479 *μ*m area of tissue across 2-3 tissue sections per animal; n=3-4 per group. Statistical significance was set to *p* < .05 with \**p* < .0332 and \*\**p* < .0021, ns=not-significant.

## Discussion

Here we investigated the use of FUS with microbubbles to induce localized BBBO for the delivery of MRI-visible albumin-based drug carriers as a method for non-invasive drug delivery to the brain. In addition, we investigated whether our albumin-based NCs were sensitive to FUS activated release of their payload to provide an element of temporal control. In order to investigate this, we custom built a benchtop FUS delivery setup capable of stereotactic MRI-guidance. We found that FUS delivered in this way was able to reliably open the BBB in two different brain regions. In addition, we were able to use FUS BBBO to localize IV injected NCs to the rat hippocampus. We were then able to induce release of both dye and glutamate from NCs with a second microbubble-free FUS treatment. Release of glutamate from NCs caused neuronal activation evident by elevated c-Fos expression, indicating that NC load was sufficient to provide localized neuromodulation. Importantly, increases in c-Fos expression were not observed in brain tissue from animals who received BBBO and release-FUS treatments without glutamate or in animals who received GluNCs without release-FUS. This is important because FUS alone (driven at different amplitudes and duty cycles than used here) has been shown to cause neuronal activation [29–32] and recently FUS BBBO alone has been reported to cause increases in BDNF, EGR1 and hippocampal neurogenesis [33].

Microbubble assisted FUS induced BBBO has gained much attention for its ability to allow localized delivery of neuro-therapeutics and neuromodulators across the BBB [38–41]. However this technique alone, requires the use of agents that do not readily cross the BBB. Combining FUS BBBO with albumin-based NCs can allow for current clinical drugs to be used in novel temporally and spatially specific ways while limiting off target effects. We loaded NCs with glutamate in order to use c-Fos expression as a marker for drug release. However, the NCs used here and other albumin-based drug carriers are able to load a wide variety of agents [17,19,34]. This use of drug encapsulation is similar to previous studies that have used FUS for triggered release from systemically circulating drug carriers [35–37]. The current study differs in that it allows the drug carrier to load into brain tissue prior to release. This ensures that when released, the drug load will immediately interact with tissue, averting the risk of the agent becoming diffused into blood. Furthermore, studies employing FUS for triggered release, rely solely on agents that readily cross the BBB. Conversely, this method can enable the research and use of drugs that do not readily cross the BBB while maintaining the benefits of encapsulation, such as location specific release and shielding from off target tissues. Thus, this technique attempts to harness the benefits of both FUS-induced drug uncaging and FUS-induced BBBO for broader drug delivery applications. Nanoparticle-based T_1_-weighted MRI contrast agents have become increasingly attractive markers for drug delivery [19,22,37,42–44]. Our recent paper demonstrates that by themselves, the ultrasmall iron oxide nanoparticles were cleared from the blood quickly through the kidneys [19,45]. However, we demonstrated that the use of cross-linked BSA increases the overall hydrodynamic radius and substantially improves circulation time to >2 hours [19]. This allowed for the accumulation of NCs in the FUS BBBO region (Figure 3). The MRI visibility of this drug carrier is particularly beneficial for in vivo studies as it can provide validation of delivery to the target region prior to drug release or drug action. Recently there has been increasing interest in the development of wearable devices for the delivery of intracranial ultrasound [46–49]. The use of such a device would be ideal for future NC drug delivery studies because it would provide the ability to activate drug release in an awake, behaving animal. Therefore, once NC delivery is confirmed with MRI, animals can recover from anesthesia, and behavior can be observed during ultrasound activated drug release. This is important because behavior studies using FUS BBBO for drug delivery are lacking due to the need for anesthesia during the FUS procedure.

Here we used NCs 100-200 nm in size; these are large compared to previous studies that successfully FUS-delivered nanoparticles in the range of 6-100 nm across the BBB [39,50–52]. Previous studies have reported the need for higher powered FUS in order to deliver larger nanoparticles across the BBB [53] which may also cause tissue damage. However, we report that IV circulating 100-200 nm NCs were able to penetrate the BBB after FUS delivered at the low power of about 0.3Mpa, and no damage was detected in T_2_-weighted MR images. This large size may also affect nanoparticle tissue migration properties [51]; at 90-minutes after delivery we did not observe any positive BSA staining or FITC-dye outside the region of EBD staining, indicating that NC location was within the region of BBBO. However, in order to fully explore NC migration more extensive time course studies must be performed. It would be ideal for NCs to remain in the BBBO region because this provides location specificity, and if larger regions of drug delivery are desired this can be achieved by targeting multiple locations with FUS BBBO.

Several limitations are present in this study. One limitation is that it is unclear how our NCs are eventually cleared from brain tissue. Albumin clearance has been shown to be mediated through glial cells [54] however, BSA in this configuration may clear differently. Furthermore, it is unclear how the other NC components such as the iron oxide NPs are cleared from brain tissue, and therefore further studies will need to be conducted to investigate the NC degradation rate and mechanism. In addition, in these studies we did not perform FUS feedback control [55] which could lead to inconsistencies in the degree of FUS BBBO. Furthermore, our benchtop approach to FUS BBBO prevented the ability to check FUS BBBO targeting before NC delivery.

## Conclusions

Here we have investigated the use of FUS BBBO mediated delivery of albumin based, MRI-visible NCs as drug carriers to achieve non-invasive, location specific neuromodulation. We report that NCs were able to load into brain tissue with FUS BBBO and provided enough MRI contrast to confirm location in target brain region. In addition, it was found that NCs were vulnerable to FUS exposure, releasing their payload *in vivo* and causing a change in their MRI contrast both *in vitro* and *in vivo*. Once NC loading was confirmed with MRI, a ‘release-FUS’ treatment was applied to cause release of either dye and or glutamate from NCs *in vivo*, providing temporal control over drug release. Importantly, glutamate release from NCs was able to cause neuronal activation, evidenced by enhanced c-Fos expression, indicating that NCs can load enough drug or neuromodulator to cause changes in neuronal activity.

## Acknowledgements

This research was supported in part by an NSF EPSCoR Research Infrastructure grant to Clemson University [NSF 1632881]. In addition, this research was supported in part by the Civitan International Research Center, Birmingham, AL. Rich M. was supported by the Alabama EPSCoR Graduate Research Scholars Award. The authors gratefully acknowledge the use of the services and facilities of the UAB Comprehensive Cancer Center’s Preclinical Imaging Shared Facility Grant [NIH P30 CA013148] and the Vision Sciences Research Center core [NIH P30 EY003039].

## Author Statement

**Megan C. Rich:** Conceptualization, Methodology, Validation, Formal Analysis, Investigation, Writing – Original Draft, Visualization. **Jennifer Sherwood:** Investigation. **Aundrea F. Bartley:** Investigation. **Quentin A. Whitsitt:** Investigation. **W. R. Willoughby:** Writing – Review & Editing. **Lynn E. Dobrunz:** Conceptualization, Writing – Review & Editing. **Yuping Bao:** Conceptualization, Resources. **Farah D. Lubin:** Conceptualization, Resources. **Mark Bolding:** Conceptualization, Methodology, Software, Investigation, Resources, Writing – Review & Editing, Supervision, Project Administration.

